# Artificial Neural Network Reveals the Role of Transport Proteins in *Rhodopseudomonas palustris* CGA009 During Lignin Breakdown Product Catabolism

**DOI:** 10.1101/2025.02.21.639544

**Authors:** Niaz Bahar Chowdhury, Mark Kathol, Nabia Shahreen, Rajib Saha

## Abstract

*Rhodopseudomonas palustris*, a versatile bacterium with diverse biotechnological applications, can effectively breakdown lignin, a complex and abundant polymer in plant biomass. This study investigates the metabolic response of *R. palustris* when catabolizing various lignin breakdown products (LBPs), including the monolignols *p*-coumaryl alcohol, coniferyl alcohol, sinapyl alcohol, *p*-coumarate, sodium ferulate, and kraft lignin. Transcriptomics and proteomics data were generated for those specific LBP breakdown conditions and used as features to train machine learning models, with growth rates as the target. Three models—Artificial Neural Networks (ANN), Random Forest (RF), and Support Vector Machine (SV)—were compared, with ANN achieving the highest predictive accuracy for both transcriptomics (94%) and proteomics (96%) datasets. Permutation feature importance analysis of the ANN models identified the top twenty genes and proteins influencing growth rates. Combining results from both transcriptomics and proteomics, eight key transport proteins were found to significantly influence the growth of *R. palustris* on LBPs. Re-training the ANN using only these eight transport proteins achieved predictive accuracies of 86% and 76% for proteomics and transcriptomics, respectively. This work highlights the potential of ANN-based models to predict growth-associated genes and proteins, shedding light on the metabolic behavior of *R. palustris* in lignin degradation under aerobic and anaerobic conditions.

**Importance:** This study is significant as it addresses the biotechnological potential of *Rhodopseudomonas palustris* in lignin degradation, a key challenge in converting plant biomass into commercially important products. By training machine learning models with transcriptomics and proteomics data, particularly Artificial Neural Networks (ANN), the work achieves high predictive accuracy for growth rates on various lignin breakdown products (LBPs). Identifying top genes and proteins influencing growth, especially eight key transport proteins, offers insights into the metabolic niche of *R. palustris*. The ability to predict growth rates using just these few proteins highlights the efficiency of ANN models in distilling complex biological systems into manageable predictive frameworks. This approach not only enhances our understanding of lignin derivative catabolism but also paves the way for optimizing *R. palustris* for sustainable bioprocessing applications, such as bioplastic production, under varying environmental conditions.

## Introduction

The climate crisis is pushing industries to adopt sustainable practices focused on efficient resource use, waste reduction, and replacing fossil fuels with renewable energy. The chemical industries traditionally rely on oil and gas due to their availability and low cost (1). However, as fossil fuel reserves deplete, with the burgeoning energy demand, renewable sources like wind, solar, geothermal, hydropower, and biomass—an abundant carbon source—are gaining much wider attention (2). Among these renewable sources, biomass conversion into useful chemicals is a promising step towards sustainability. Of the 170 billion tons of renewable resources produced annually, only 3.5% is utilized by humans (3). Lignin, a major component of lignocellulosic biomass, is increasingly seen as a valuable resource for producing fuels and chemicals (4). Found in plant cell walls, lignin provides structural support and is abundant in wood and agricultural residues. Its complex molecular structure, comprising three phenylpropane units linked by ether and carbon–carbon bonds, along with hydroxyl and methoxy groups, makes it highly resistant to degradation (5). This complexity presents a challenge in efficiently converting lignin into high-value products (6). To overcome that, microbial degradation of lignin into desirable products can be an efficient alternate route.

*Rhodopseudomonas palustris* CGA009 is an alphaproteobacterium capable of thriving in diverse metabolic environments, including both phototrophic and chemotrophic conditions. It has the remarkable ability to fix carbon dioxide and nitrogen and can grow under either aerobic or anaerobic conditions. This versatility allows it to produce ATP using light, as well as organic or inorganic compounds for energy generation (7, 8). *R. palustris* was previously shown to catabolize *p*-coumarate, a lignin breakdown product (LBP) (9). Thereby it can be a promising bacterium to degrade different LBPs into value-added chemicals. However, to achieve that goal, a system-wide understanding of *R. palustris* is essential. A system-wide understanding of *R. palustris* is required to effectively apply it for converting lignin breakdown products (LBPs) into value-added chemicals because it allows us to identify and optimize the complex metabolic and regulatory process involved in LBP catabolism.

Genome-scale metabolic models (GSMs) are popular tools to gain systems-wide understanding of a living system (10, 11). So far, two GSMs of *R. palustris* were reconstructed. The earlier GSM of *R. palustris* (12) was used to quantitatively understand trade-offs among a set of important biological objectives during different metabolic growth modes. Later, we developed another updated GSM of *R. palustris* (iRpa940), with an 84% accuracy, that successfully captured the relation between light-dependent energy production and oxidation rate of quinol (13). Moreover, based on iRpa940, we reconstructed the first-ever Genome-scale metabolic and expression (ME-) model of *R. palustris* which successfully captured ferredoxin as a regulatory element in distributing electrons between carbon fixation and nitrogen fixation pathways, two major redox balancing pathways in *R. palustris* (14). However, these first principle models usually connect metabolic reactions with bacterial growth rate. Thus, quantifying the relationship between non-metabolic genes/proteins and bacterial growth rate is not intuitive, as seen with enzymes like *boxB*, involved in anaerobic aromatic compound degradation, and *nuoF2*, which functions in the electron transport chain. In contrast, non-metabolic genes/proteins participate in processes like transport, signaling, stress response, and regulation, as exemplified by rpa2624 (a sulfonate transport protein). However, conventional techniques such as differential expression analysis and pathway enrichment typically focus on individual genes or proteins with significant expression changes, potentially missing subtle, non-linear interactions and the combined effects of multiple factors. To resolve this issue, machine learning tools can be implemented. Previously, machine learning (ML) tool was used to identify proteins that are associated with biomarkers in COVID-19 (15). Moreover, ML was used to successfully identify proteins in hepatic carcinoma (16). Furthermore, ML was used to predict antimicrobial proteins for numerous bacterial species (17). Therefore, despite being a black box approach, ML can be a promising tool to identify proteins, metabolic and/or non-metabolic, associated with bacterial growth rate.

In this work, we generated growth profiles, transcriptomics, and proteomics data for *R. palustris* as it catabolized various lignin breakdown products (LBPs), including monolignols (*p*-coumaryl alcohol, coniferyl alcohol, sinapyl alcohol), acid derivatives (p-coumarate and sodium ferulate), and kraft lignin and investigated their catabolic routes, with results included in a companion paper (18). Using these transcriptomics and proteomics datasets as features and growth rates as targets, three ML models—Artificial Neural Networks (ANN), Random Forest (RF), and Support Vector Machine (SVM)—were trained, with ANN achieving the highest accuracy (94% for transcriptomics and 96% for proteomics). Since we wanted to pick genes and proteins which are directly associated with growth, we used the entire data set to train the ML models using standard

protocol (15). Next, permutation feature importance analysis on the ANN models revealed the top twenty genes and proteins most impacting growth, and combining these insights identified eight key transport proteins (mostly associated with amino acid and sulfonate transportation) driving *R. palustris* growth on LBPs. Retraining the ANN using these eight proteins yielded 86% accuracy for proteomics data and 76% for transcriptomics data. This work highlights the effectiveness of ANN models in predicting growth-associated genes and proteins, offering broader implications for optimizing microbial systems in bioconversion processes.

## Results and Discussion

### Growth of *R. palustris* on different lignin breakdown products

Although the ability of *R. palustris* to catabolize *p*-coumarate is already established (9), in this work, we wanted to reassess its capability to catabolize other LBPs as well. Wild-type cultures were supplemented with various LBPs, including kraft lignin. Specifically, the monolignols *p*-coumaryl alcohol, coniferyl alcohol, and sinapyl alcohol, along with two acid derivatives, *p*-coumarate and sodium ferulate, were evaluated as potential carbon sources. These substrates were selected based on their prevalence in the depolymerization of kraft lignin. Acetate, a carbon source with a simple catabolic route (13), was included as a positive control. Growth curves were generated for each substrate under both aerobic and anaerobic conditions (Supplementary Figure S1), and growth rates were calculated using logistics model.

Notably, *R. palustris* is unable to grow on certain LBPs when these are the sole carbon sources under varying oxic conditions, corroborating findings from previous studies (19, 20). In such cases, 10 mM of sodium acetate was supplemented as an additional carbon source. However, to confirm that *R. palustris* is utilizing the LBPs for biomass production rather than relying on acetate, the OD660 of these cultures must have exceeded that of the positive acetate control. Among the substrates tested, only *p*-coumarate was consistently consumed by *R. palustris* without the need for acetate supplementation, under both aerobic and anaerobic conditions. In contrast, *p*-coumaryl alcohol and the methoxylated monomers were more resistant to degradation. We hypothesize that acetate, which directly enters the TCA cycle, efficiently generates ATP to support the energy-intensive catabolism of lignin breakdown products (LBPs), providing the necessary energy to drive their enzymatic degradation.

Building on the initial experiments, which provided clear evidence of *R. palustris’* ability to catabolize these LBPs, we further explored the organism’s transcriptomic and proteomic response when metabolizing these substrates. For transcriptomic analysis, two biological replicates were harvested during the mid-exponential growth phase (Supplementary Table S1) and submitted for Next-Generation Sequencing (NGS). Additionally, five biological replicates for each condition were collected to analyze the proteome profiles. We also wanted to make sure that photosynthetic machinery was active in the anoxic growth. One key regulator, *fixK*, was proportionally upregulated in our anaerobic samples compared to aerobic samples, suggesting its role in adapting to anoxic conditions. Additionally, we observed reduced abundance of genes encoding oxygen-dependent enzymes such as *hemF* (RPA1514), cytochrome bd oxidases (RPA1319, RPA4452, RPA4793-RPA4794), and cytochrome aa3 oxidases (RPA1453, RPA4183, RPA0831-RPA0836) (Table S4). In contrast, genes associated with high-affinity cytochrome cbb3 oxidase (RPA0015-RPA0019), oxygen-independent coproporphyrinogen oxidase HemN (RPA1666), and components of the photosynthetic apparatus (RPA1505-RPA1507, RPA1521-RPA1548, RPA1667-RPA1668, RPA3568) were upregulated. These findings indicate a strong activation of photosynthetic machinery and related pathways under anoxic, light-exposed conditions, consistent with *R. palustris* physiology. This system-wide shift in metabolism, redox balance, and pigment biosynthesis confirms that our photoheterotrophic cultures were indeed anaerobic, as also validated by *FixK*-regulated gene induction (RPA1006-RPA1007, RPA1554). These observations inform our analysis by demonstrating how anaerobic growth in the presence of light drives specific metabolic and regulatory adaptations. By training different machine learning (ML) models where transcriptomics/proteomics data will be feature and the growth rates will be target, we can identify important genes/proteins associated with the *R. palustris* growth under aerobic and anaerobic conditions for different LBPs. The overall workflow of the work can be found in Fig. 1a and b.

**Fig 1:**
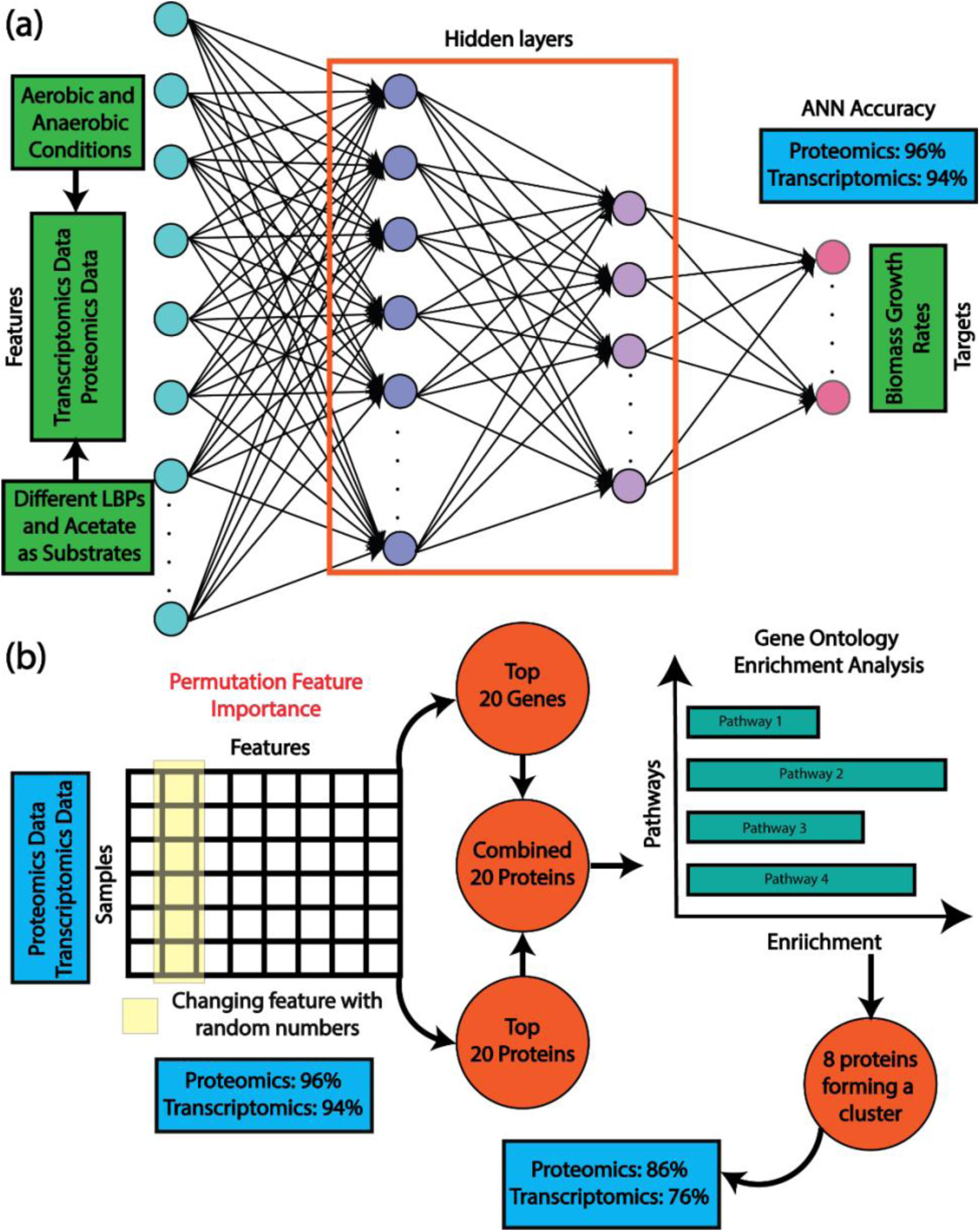
Artificial Neural Network (ANN) predicts top 20 growth-associated genes and proteins. (a) ANN predicted growth rates from transcriptomics and proteomics data with very high accuracy. (b) Later permutation feature importance was used to determine the top twenty genes and proteins, affecting the growth rates most. Furthermore, a list of top twenty proteins, determined from the combined list of top twenty genes and top twenty proteins, yielded eight proteins through gene set enrichment analysis and. Further training the ANN only with those eight proteins and genes resulted in 86% and 76% accuracy respectively.

### Artificial Neural Networks accurately predicted growth rates from ‘omics’ data

Once the transcriptomics and proteomics data were generated, these datasets were used to build different ML algorithms. As proteins are often difficult to detect compared to the mRNA, for training ML algorithms, we only kept the genes in the transcriptomics dataset that matched with detected proteins. Furthermore, we normalized both proteomics and transcriptomics dataset by subtracting mean and scaling it to the unit variance. In the process, we were able to keep 1855 genes and 1855 proteins to train ML algorithms for fourteen different conditions.

Next, for both the transcriptomics and proteomics dataset, we benchmarked three different machine learning models – artificial neural network (ANN), random forest algorithm (RF), and support vector machine (SV) to assess the suitability of each to predict growth rates based on transcriptomics and proteomics data individually. In these algorithms, proteomics/transcriptomics data were the features and growth rates were the target. As the aim of this study is to pin-point which proteins/genes have the most impact on the growth of *R. palustris* for LBP catabolism, for all the algorithms, following the previously published work (15), we used the entire datasets (transcriptomics/proteomics) to train each model. In that way, the entire genome was used to train the ML models against growth rates, which is more realistic than using only a part of the genome to train ML models. Among these three algorithms, ANN achieved the least mean absolute error (MAE) for both transcriptomics and proteomics data (0.00129 and 0.00111 respectively), followed by RF (0.00547 and 0.00524 respectively) and SV (0.00859 and 0.00841) (Fig. 2a). Similarly, ANN achieved the least mean square error (MSE) for both transcriptomics and proteomics data (2.76 × 10^−6^ and 2.19 × 10^−6^ respectively), followed by RF (3.69 × 10^−5^ and 3.45 × 10^−5^ respectively) and SV (8.23 × 10^−5^ and 7.92 × 10^−5^) (Fig. 2b). In terms of accuracy, ANN showed much higher accuracy in predicting the growth rates from transcriptomics and proteomics data (93.68% and 95.99% respectively) compared to the RF (76.28% and 77.29% respectively) and SV (65.89% and 66.32% respectively) (Fig. 2c).

**Fig 2:**
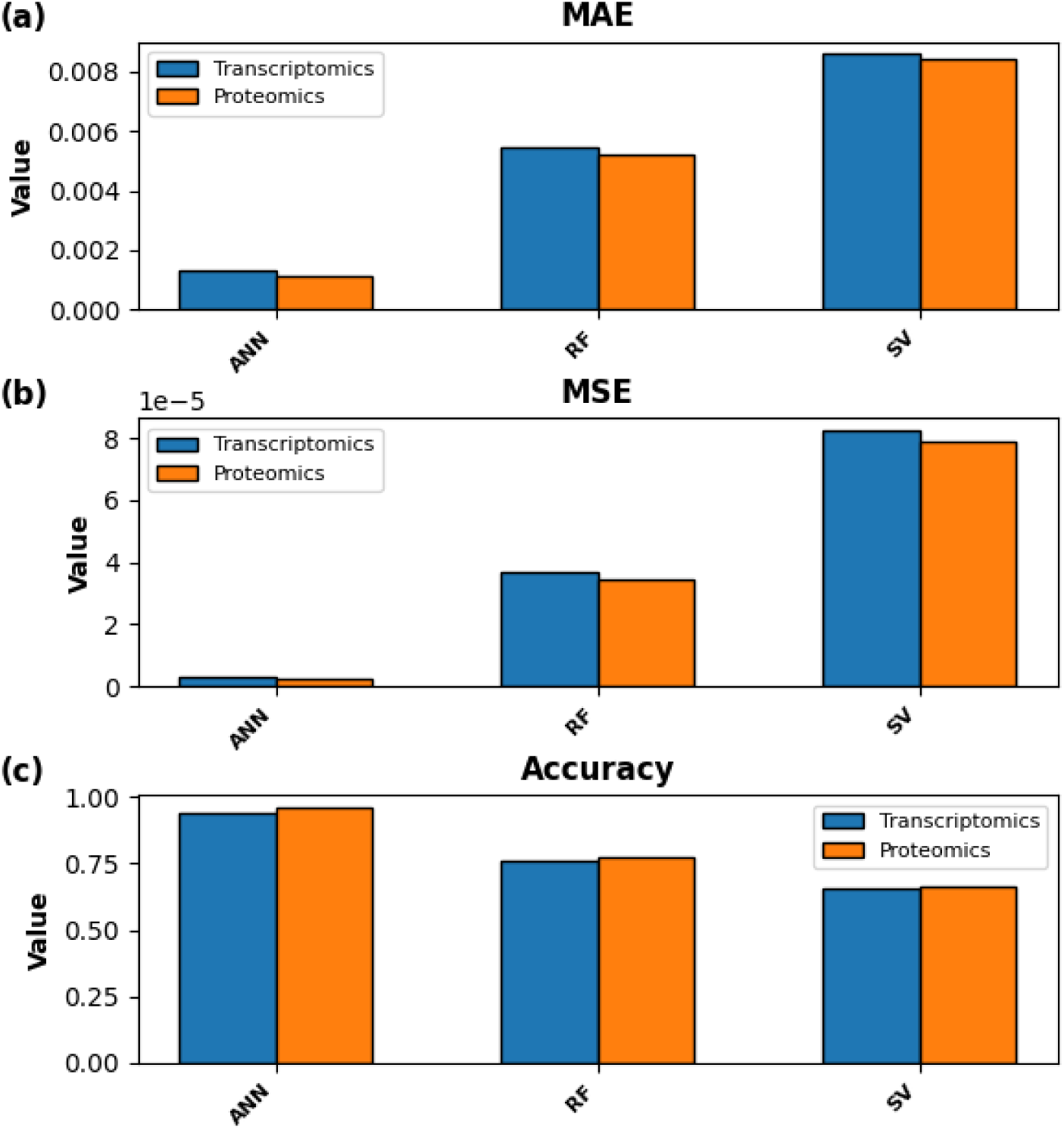
Artificial Neural Network (ANN) is the best performing machine learning algorithm compared to Random Forest (RF) algorithm and Support Vector (SV) Machine in *R. palustris*. (a) Mean Absolute Error (MAE) for three different machine learning algorithms. (b) Mean Squared Error (MSE) for all three different machine learning algorithms. (c) Accuracy for three different machine learning algorithms.

Overall, from the training of three different machine learning algorithm, ANN was the best performing algorithm to capture growth rates from transcriptomics and proteomics data. Moreover, proteomics data is a better indicator of enzymatic activity compared to the mRNA, as the latter one undergoes fast dilution and degradation due to its unstable nature (21).

### Permutation Feature Importance identified top twenty growth-associated genes

As ANN was the best performing machine learning algorithm, we choose ANN to find the most important genes and proteins that impacted the growth rates. At first, from the ANN algorithm, we generated top twenty genes impacting the growth rates using permutation feature importance from all the given conditions (22) (Fig. 3a). Permutation feature importance is an algorithm used to evaluate the importance of features in a ML model by measuring how performance decreases when a feature’s values are randomly shuffled. The underlying idea is that if a feature is important for making accurate predictions, randomly shuffling its values should lead to a significant drop in model performance. If shuffling a feature does not affect performance, the model doesn’t rely on that feature to make predictions. The expression pattern of those twenty genes can be observed in Fig. 2b.

**Fig 3:**
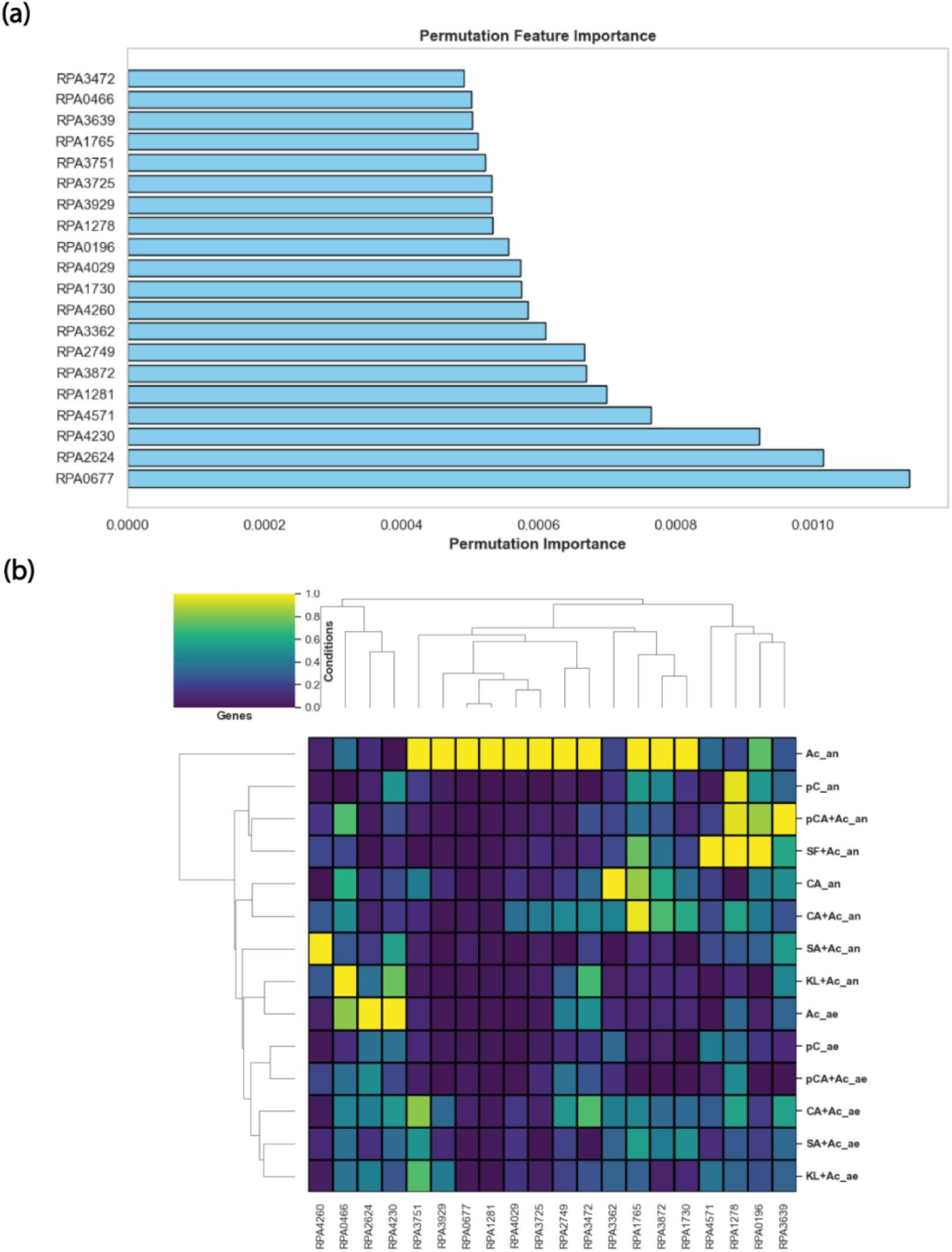
ANN predicted the top twenty genes that affected the growth rate most. (a) Top twenty growth associated genes from the deep learning framework using permutation feature importance. (b) Top twenty growth associated gene expression profile.

From twenty top genes (Supplementary Table S2), there were only four genes that were directly associated with metabolism. *rpa0677* (*boxB*), *rpa4260* (*nuoF2*), *rpa1765* (*ech*), and *rpa3472* (*ilvD1*). We determined the metabolic activity of a given gene based on KEGG database. For example, if a gene is associated with a metabolic reaction, that gene is considered to be directly associated with metabolism. Benzoyl-CoA 2,3-epoxidase (*boxB*) plays a key role in the anaerobic degradation of aromatic compounds, particularly benzoate, in *R*. *palustris*. This enzyme is part of the pathway that enables *R. palustris* to utilize aromatic compounds as a carbon and energy source under anaerobic conditions. *boxB* catalyzes the epoxidation of benzoyl-CoA, forming 2,3-epoxybenzoyl-CoA, which is a crucial step in the ring cleavage and degradation of the aromatic structure thereby may play a role in degradation of many other LBPs (23). Moreover, *boxB* often works with a redox partner (e.g., flavoprotein reductase) to receive electrons and enable the epoxidation reaction, utilizing reduced electron carriers such as NADPH or ferredoxin (24). Next, NADH-quinone oxidoreductase (*nuoF2*) in *R. palustris* plays a key role in its electron transport chain by transferring electrons from NADH to quinones, such as ubiquinone, while simultaneously pumping protons across the membrane. This creates a proton gradient used to generate ATP, crucial for the cell’s energy production. In *R. palustris*, which thrives in diverse metabolic conditions (8), *nuoF2* helps maintain redox balance, adapting to various electron acceptors and energy sources (25). Enoyl-CoA hydratase (*ech*) plays a key role in the β-oxidation of fatty acids, catalyzing the hydration of enoyl-CoA to 3-hydroxyacyl-CoA. This is essential for fatty acid degradation, providing energy and carbon for the cell; additionally, it may be involved in the degradation of hydrocarbons, helping to metabolize alkenes and alkanes (26). Finally, xylonate dehydratase (*xylD*) plays a crucial role in the metabolism of pentose sugars, specifically in the conversion of xylonate to 2-keto-3-deoxyxylonate, a key intermediate in the pentose and glucuronate interconversion pathways (27). While *R. palustris* cannot directly catabolize xylose or glucose, this enzyme may facilitate the utilization of related metabolites derived from plant-derived polysaccharides, supporting its adaptation to lignocellulose-rich environments.

Next, from gene ontology analysis of those twenty genes using ShinyGO (28), we found only one cluster of eight genes, associated with signalling and transportation mechanism of different substrates. These genes are *rpa0466* (metalloprotease inhibitor), *rpa2624* (sulfonate transport), *rpa3362* (hypothetical protein), *rpa3725* (ABC transporter substrate-binding protein), *rpa3751* (cache domain-containing protein), *rpa3872* (hypothetical protein), *rpa4029* (ABC transporter substrate-binding protein), and *rpa4571* (BA14K family protein). In *R. palustris*, the listed genes represent a diverse array of functions that contribute to the organism’s metabolic versatility and environmental adaptability. *rpa0466*, a metalloprotease inhibitor, likely plays a regulatory role in proteolysis, protecting proteins from degradation under certain conditions (29). *rpa2624* is involved in sulfonate transport, enabling the organism to utilize sulfur from organic sources, which is important in sulfur-limited environments (30). Both *rpa3725* and *rpa4029* are substrate-binding proteins of ABC transporters, which facilitate the uptake of various substrates critical for nutrient acquisition (31). *rpa3751*, a cache domain-containing protein, is likely involved in sensing extracellular signals or nutrients, guiding cellular responses (32). *rpa3362* and *rpa3872*, both hypothetical proteins, might represent uncharacterized functions in metabolism or stress response. Lastly, *rpa4571*, a BA14K family protein, could have a role in stress resistance or environmental interactions (33). Together, these genes reflect the complex regulatory, transport, and environmental sensing capabilities of *R. palustris*, enhancing its ability to thrive in diverse ecosystems. Comparing the result with gene ontology analysis, StringDB database could not predict any interaction among the proteins encoded by these genes. The lack of predicted interactions in StringDB, along with the high prevalence of hypothetical and unannotated genes in *R. palustris*, underscores the unique and relatively understudied nature of this organism.

The rest of the genes, such as *rpa4230*, *rpa1281*, *rpa2749*, *rpa1730*, *rpa0196*, *rpa1278*, *rpa3929*, *rpa3639* did not fall into specific categories in gene ontology analysis with *rpa4230* was identified as gene with hypothetical function. Details of rest of the genes can be accessed in the Supplementary Table S2.

### Permutation Feature Importance identified top twenty growth-associated proteins

Similar to the twenty top genes, we also identified top twenty growth-associated proteins from all the given conditions (Supplementary Table S3) using permutation feature importance from the ANN model trained using proteomics data (Fig. 4a). Their abundance profile can be observed in Fig. 4b. Interestingly, there is no overlap between the list of top twenty genes and top twenty proteins, thus indicating weak correlation between transcriptomics and proteomics data. The lack of overlap between transcriptome and proteome datasets indicates a weak correlation between mRNA and protein levels, a common phenomenon in alphaproteobacteria like *Rhodobacter sphaeroides* (34). This suggests significant post-transcriptional regulation, where factors such as mRNA stability, translation efficiency, and protein turnover decouple transcript and protein abundances. Among those proteins, three were associated with metabolism: RPA0069 (tryptophan synthase subunit beta), RPA0532 (beta-ketoacyl-ACP reductase), and RPA2720 (glutathione-dependent disulfide-bond oxidoreductase). In *R. palustris*, the tryptophan synthase subunit beta plays a vital role in the biosynthesis of the essential amino acid tryptophan. This enzyme catalyzes the final step in the tryptophan biosynthesis pathway, converting indole and serine into tryptophan (35). As tryptophan is a precursor for several important biomolecules, including proteins and signaling compounds, its synthesis is critical for *R. palustris* to maintain cellular function and adapt to various environmental conditions. The ability to synthesize tryptophan internally allows *R. palustris* to thrive in nutrient-limited environments where external sources of amino acids may be scarce, contributing to its metabolic independence and ecological versatility. Beta-ketoacyl-ACP reductase is a key enzyme in the fatty acid biosynthesis pathway. It catalyzes the reduction of beta-ketoacyl-ACP to beta-hydroxyacyl-ACP, an essential step in the elongation cycle of fatty acid synthesis. These fatty acids are crucial components of the cell membrane and are also used for energy storage. By facilitating the production of lipids, beta-ketoacyl-ACP reductase plays a critical role in maintaining membrane integrity, enabling adaptation to environmental stress, and supporting energy metabolism (36). This enzyme’s function is fundamental to the organism’s ability to build cellular structures and thrive in diverse ecological niches. Finally, glutathione-dependent disulfide-bond oxidoreductase plays a crucial role in maintaining cellular redox balance by catalyzing the reduction of disulfide bonds in proteins, using glutathione as a cofactor (37). This enzyme is essential for protecting the cell against oxidative stress by facilitating proper protein folding, repair, and regeneration under adverse conditions. By reducing oxidized protein thiols, it helps mitigate damage caused by reactive oxygen species, thereby preserving protein function and structural integrity. Additionally, it supports broader cellular defense mechanisms against environmental fluctuations, enhancing the organism’s adaptability. By maintaining redox homeostasis, glutathione-dependent disulfide-bond oxidoreductase enables *R. palustris* to efficiently manage oxidative stress, sustain metabolic functions, and thrive in dynamic environments where redox imbalances could compromise cell viability.

**Fig 4:**
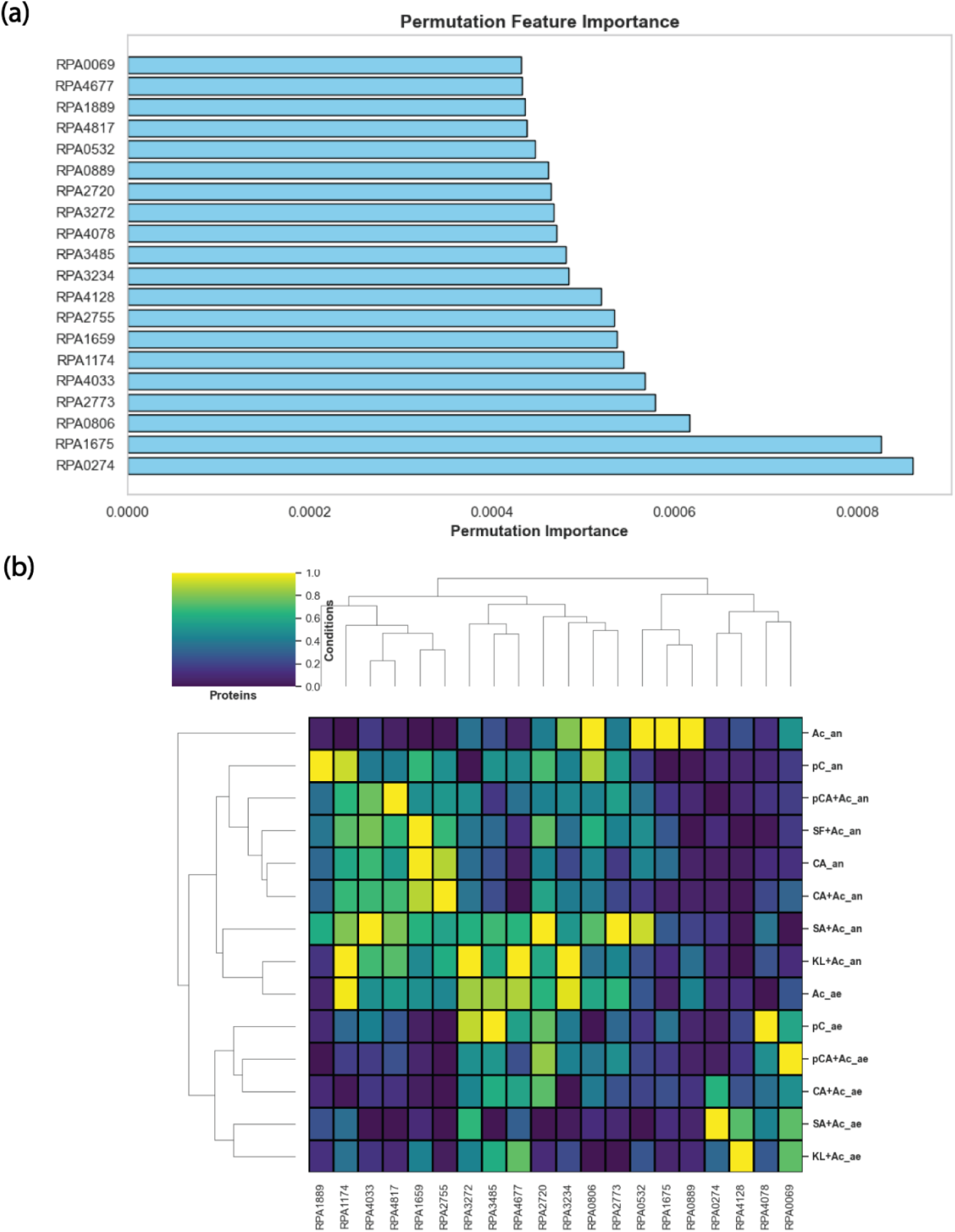
ANN predicted the top twenty proteins that affected the growth rate most. (a) Top twenty growth associated proteins from the deep learning framework using permutation feature importance. (b) Top twenty growth associated proteins abundance profile.

Next, the StringDB network reveals that only RPA3272 (large subunit ribosomal protein L1) and RPA3234 (50S ribosomal protein L18), out of the other proteins, were strongly connected with each other among those twenty proteins (Supplementary Figure S2). Besides, a previous gene essentiality study found both proteins as essential (38). In *R. palustris*, RPA3272 and RPA3234 are essential components of the ribosome, which plays a critical role in protein synthesis. RPA3272 is involved in binding rRNA and is important for the release of tRNA during translation, ensuring efficient protein production (39). RPA3234 helps stabilize the structure of the 50S ribosomal subunit by binding to 5S rRNA, which is crucial for the assembly and function of the ribosome (39). These ribosomal proteins ensure the accurate and efficient synthesis of enzymes and proteins that drive key metabolic pathways, including photosynthesis, nitrogen fixation, and carbon utilization. By supporting robust protein translation, RPA3272 and RPA3234 contribute to the metabolic flexibility and environmental adaptability of *R. palustris*.

While the rest of the proteins, such as RPA4677, RPA1889, RPA4817, RPA0889, RPA4078, RPA3485, RPA4128, RPA2755, RPA1659, RPA1174, RPA4033, RPA0806, RPA1675, and RPA0274 did not fall into specific categories in gene ontology/StringDB analysis, we explored their role on an individual basis. The caspase family protein (RPA4677) could be involved in programmed cell death or stress responses, playing a role in cellular regulation (40). Domain-containing proteins such as DUF2019 (RPA1889) and DUF2188 (RPA4128) suggest the presence of proteins with unidentified functions, potentially linked to regulatory or structural roles within the cell. The response regulator transcription factor (RPA4817) may participate in signal transduction pathways, enabling *R. palustris* to respond to environmental stimuli (41). Proteins like the molecular chaperone (RPA0889) and DNA starvation/stationary phase protection protein (RPA2755) are likely involved in stress protection, helping the bacterium survive adverse conditions. Enzymes such as CoA transferase (RPA3485), uroporphyrinogen-III synthase (RPA4033), and 23S rRNA methyltransferase (RPA1174) point to key roles in metabolic pathways and gene expression. Proteins related to transport and chemotaxis, such as the ABC transporter substrate-binding protein (RPA0806), the methyl-accepting chemotaxis protein (RPA1675), and the P-II family nitrogen regulator (RPA0274), are likely crucial for nutrient uptake and environmental navigation, reflecting the bacterium’s ability to thrive in diverse habitats. Overall, these proteins highlight the complex regulatory and metabolic network that underpins the survival and versatility of *R. palustris*. Details of rest of the proteins can be accessed in the Supplementary Table S3.

### Statistical analysis revealed regulatory and conserved proteins during lignin breakdown

Since these top twenty proteins have diverse functions, we also wanted to investigate how their functionality varies with the availability of oxygen. Thus, we determined how many of the top twenty most important proteins showed statistically significant differences in abundance between aerobic and anaerobic conditions. Of these top twenty proteins, we found that nine exhibited statistically significant differences: RPA0069, RPA1889, RPA4817, RPA4078, RPA4128, RPA2755, RPA1659, RPA1174, and RPA4033 (Fig. 5).

**Fig 5:**
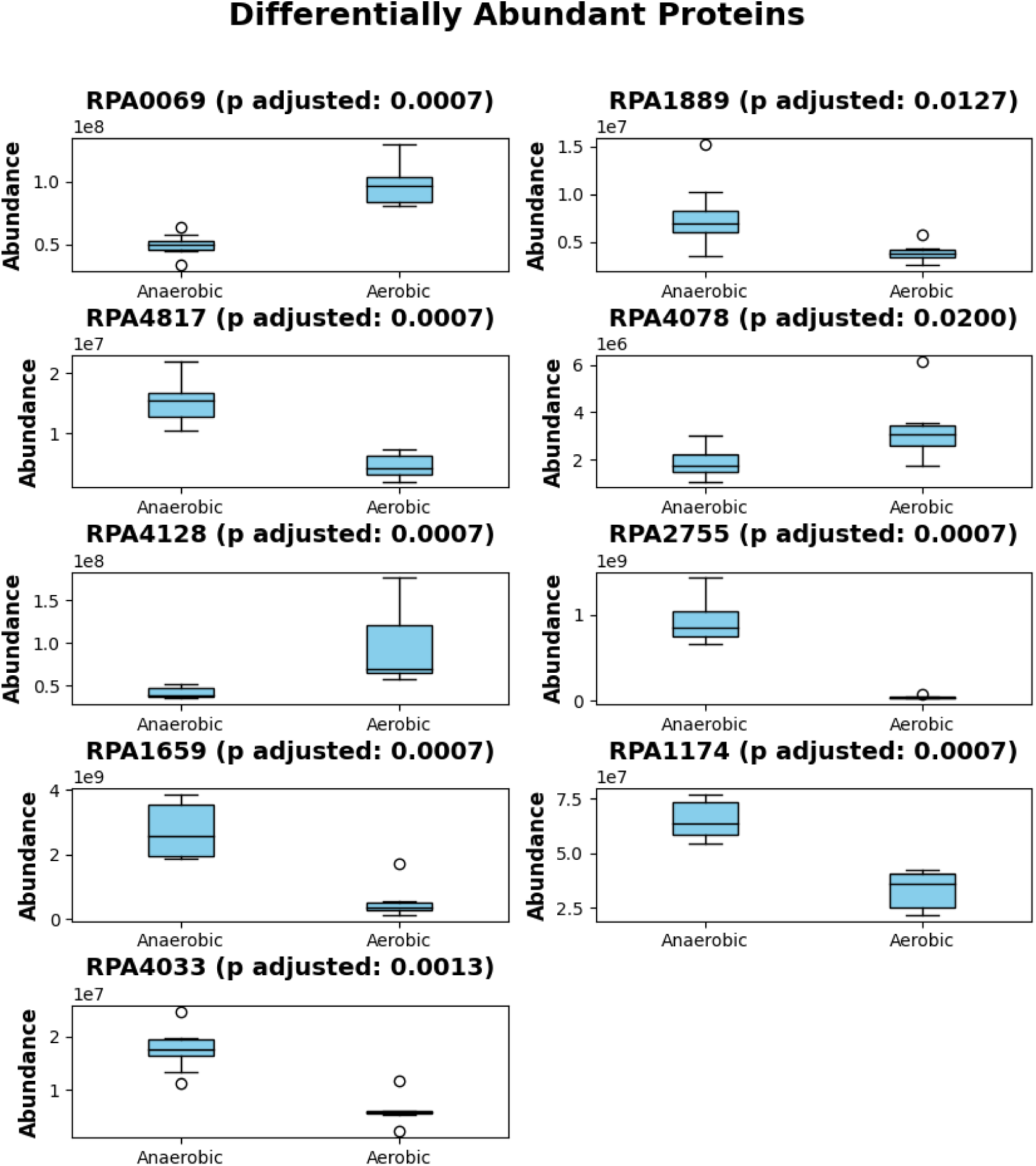
Out of top twenty growth associated proteins, nine are associated with aerobic conditions.

The differential abundance of these proteins between aerobic and anaerobic conditions suggests that these proteins play distinct roles in cellular responses to oxygen availability. For instance, tryptophan synthase subunit beta (RPA0069) may be involved in metabolic pathways that are more active under oxygen-rich environments, where biosynthetic processes like amino acid synthesis require higher energy. The response regulator transcription factor (RPA4817) could indicate alterations in gene regulation in response to changes in redox conditions. Proteins such as the DNA starvation/stationary phase protection protein (RPA2755) and hemerythrin domain-containing protein (RPA1659) are likely involved in the cell’s adaptive response to stress, where the lack of oxygen imposes survival challenges, prompting the activation of protective or metabolic adjustments. The presence of DUF domain-containing proteins (e.g., RPA1889 and RPA4128) which have unknown or poorly characterized functions, might represent novel pathways or processes important in anaerobic adaptation. Moreover, the activity of enzymes such as 23S rRNA methyltransferase (RPA1174) and uroporphyrinogen-III synthase (RPA4033) hints at modifications in translation machinery and heme biosynthesis, respectively, both of which are crucial under varying oxygen levels, potentially impacting cellular respiration and energy production. Finally, the appearance of a hypothetical protein (RPA4078) highlights that there might still be unknown factors contributing to the organism’s response to oxygen tension.

Among those nine proteins, whose abundances changed significantly between aerobic and anaerobic conditions, we also wanted to evaluate which proteins had the highest differences in abundances. To accomplish that, we calculated the coefficient of variation (CV) for all the nine proteins across aerobic and anaerobic conditions (Supplementary Figure S3). The higher the CV is, the more differentially abundant the protein is between aerobic and anaerobic conditions. From the CV analysis, RPA2755 (DNA starvation/stationary phase protection protein) showed the highest CV between aerobic and anaerobic conditions. RPA1174 (23S rRNA methyltransferase) showed the least CV among those nine proteins.

The rest of the 11 (RPA4677, RPA0532, RPA0889, RPA2720, RPA3272, RPA3485, RPA3234, RPA2773, RPA0806, RPA1675, and RPA0274) out of those top twenty proteins did not exhibit statistically significant differences in their abundances between aerobic and anaerobic conditions. This lack of differential abundance suggests that their roles are not strongly influenced by the availability of oxygen. For example, RPA4677, a caspase family protein, may participate in processes like programmed cell death, but oxygen levels do not appear to impact its regulation. Similarly, RPA0532, a beta-ketoacyl-ACP reductase involved in fatty acid synthesis, and RPA0889, a molecular chaperone, which assists in protein folding, remain unaffected by shifts in oxygen levels. Even proteins with roles in redox balance, such as RPA2720, a glutathione-dependent disulfide-bond oxidoreductase, and RPA3485, a CoA transferase linked to metabolic processes, show no significant response to oxygen conditions. Ribosomal proteins like RPA3272 and RPA3234, essential for protein synthesis, as well as RPA0806, an ABC transporter substrate-binding protein, and RPA1675, a methyl-accepting chemotaxis protein, are similarly unaffected. The hypothetical protein RPA2773 and the P-II family nitrogen regulator RPA0274 also remain stable regardless of oxygen availability. This suggests that while these proteins play essential roles in the cell, their expression is not directly modulated by aerobic or anaerobic growth, indicating their functions may be more constant across different oxygen environments.

### Combined Permutation Feature Importance highlights amino acid transporters’ role in LBP

As encoding proteins is a process starting from the mRNA, we thereby combined the permutation feature importance score for both the transcriptomics and proteomics dataset and came up with a combined permutation feature importance for each of the proteins. The list can be accessed in the Supplementary Table S4. Form the list, similar to the previous section, we identified top twenty proteins. From the list of these twenty proteins, five came from the list of top twenty proteins (RPA4677, RPA0806, RPA1675, RPA0274, and RPA0532) and ten came from the list of top twenty genes (RPA2624, RPA4571, RPA3362, RPA3872, RPA0677, RPA4230, RPA0196, RPA1281, RPA2749, and RPA3929). Interestingly, five came from outside the list of top twenty proteins and genes (RPA4797, RPA1428, RPA2455, RPA1190, and RPA4297), showcasing the importance of combining transcriptomics and proteomics data to predict the growth rates of *R. palustris* under different LBPs.

Among those five proteins, which came outside of the list of top twenty proteins and genes, 7-carboxy-7-deazaguanine synthase (RPA1190) and aldo/keto reductase (RPA4297) were the metabolic proteins. In *R. palustris*, both RPA1190 and RPA4297 play important roles in catabolizing LBPs under both aerobic and anaerobic conditions. RPA1190 is involved in the biosynthesis of 7-deazapurines, which can influence metabolic pathways associated with lignin-derived aromatic compound processing (42). Its activity might be crucial for cofactor biosynthesis that aids in the degradation of complex lignin structures. On the other hand, RPA4297, an aldo/keto reductase, catalyzes the reduction of aldehydes and ketones, which are common intermediates in lignin depolymerization. Under aerobic conditions, it helps detoxify reactive oxygen species generated during lignin degradation, while in anaerobic conditions, it assists in reducing aromatic aldehyde intermediates to less toxic alcohols, facilitating the complete mineralization of lignin-derived compounds (43). Three other proteins were also predicted outside of the top twenty list of genes and proteins. Proteins such as the amino acid ABC transporter substrate-binding protein (RPA4797), the MetQ/NlpA family ABC transporter substrate-binding protein (RPA1428), and the M23 family metallopeptidase (RPA2455) are vital for nutrient acquisition and environmental adaptation during lignin breakdown. RPA4797 plays a key role in importing amino acids, which are crucial for cellular growth and enzyme production necessary for lignin degradation pathways (44). Similarly, RPA1428, a member of the MetQ/NlpA family protein, is involved in binding substrates such as sulfur-containing compounds like methionine, which can be essential for cellular redox (45) balance during lignin depolymerization. RPA2455, a metallopeptidase from the M23 family, facilitates the breakdown of peptide bonds in proteins or small peptides that may accumulate from microbial community interactions or host biomass degradation (29). By processing and recycling these molecules, these proteins help *R. palustris* efficiently metabolize lignin-derived compounds and thrive under varying environmental conditions, optimizing nutrient acquisition and waste processing. Together, these proteins contribute to the efficient utilization of lignin as a carbon source, allowing *R. palustris* to thrive in diverse environments.

Next, we again performed gene ontology analysis using ShinyGO and protein-protein interaction using StringDB. StringDB returns no interactions among all these proteins. However, gene ontology analysis identified eight amino acid transport proteins (RPA4797, RPA1428, RPA4677, RPA0806, RPA2624, RPA4571, RPA3362, and RPA3872), associated with signalling, forms a cluster (Fig. 6a). Furthermore, only these eight proteins were used to train the ANN algorithm again, which resulted in 86% accuracy (Fig. 6b). We also used corresponding eight genes to train the ANN algorithm again, which resulted in 78% accuracy (Fig. 6b). Similar to the previous cases, proteomics data-based predictions of growth rate performed better compared to the transcriptomics data-based growth predictions using ANN. Permutation feature importance again ranked these transport proteins from the most important to the least. RPA3362, a hypothetical protein, was found to be the most important protein dictating the growth and RPA4677, a caspase protein, was identified to be the least important among those eight proteins (Fig. 6c). Thus, by combining permutation feature importance for both transcriptomics and proteomics dataset, we identified eight proteins which were able to predict the growth rates of *R. palustris* with an excellent accuracy.

**Fig 6:**
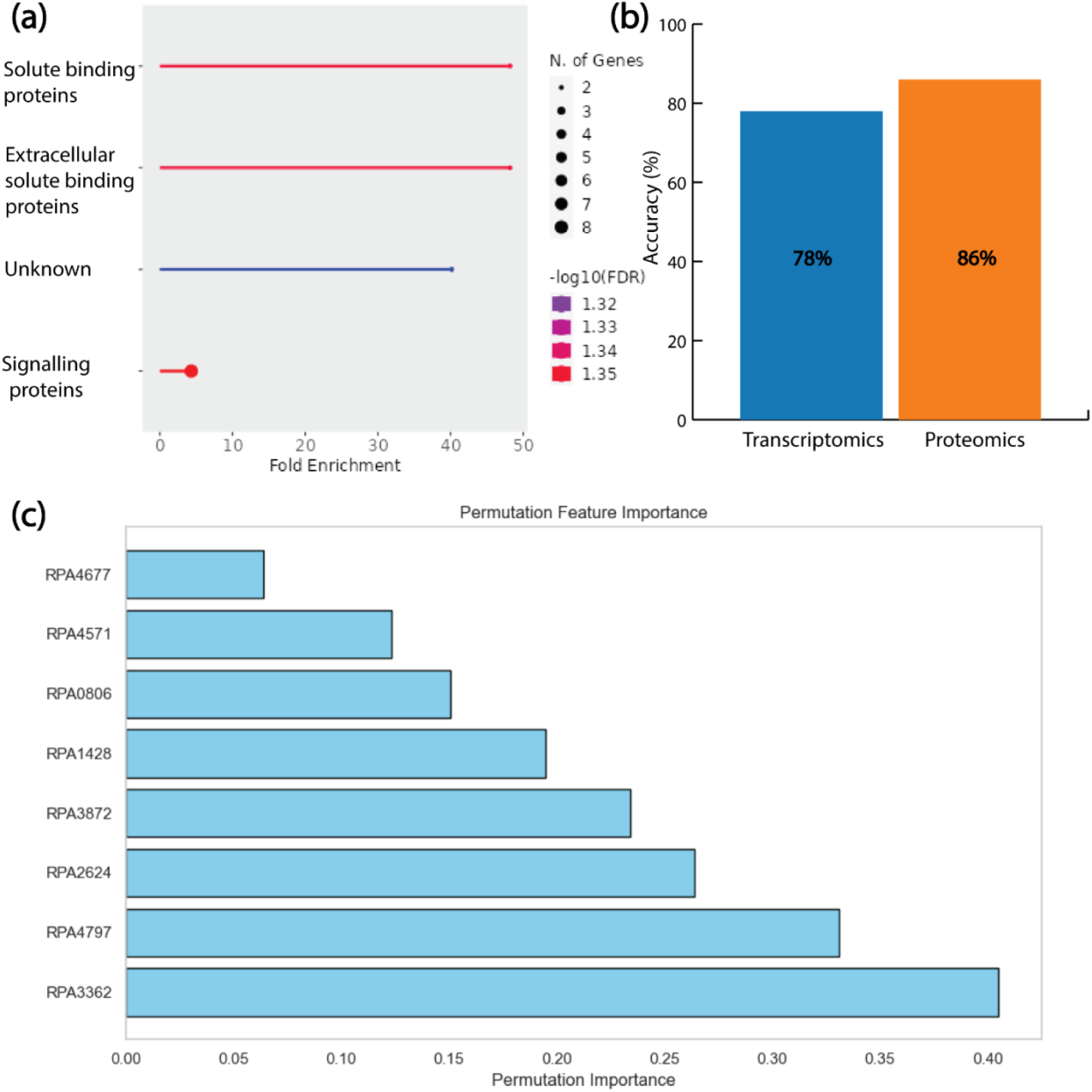
Only amino acid transport proteins can explain the growth pattern of *R. palustris* for LBPs. (a) Gene Ontology analysis revealed the role of transport protein as the signaling molecules. (b) Eight transport proteins can capture growth rate in different conditions with 86% accuracy compared to the transcriptomics data’s 78% accuracy. (c) Permutation feature importance revealed top transport proteins.

This study generated growth profiles along with transcriptomics and proteomics dataset for *R. palustris* during the catabolism of various lignin breakdown products (LBPs), including monolignols (p-coumaryl alcohol, coniferyl alcohol, sinapyl alcohol), acid derivatives (p-coumarate, sodium ferulate), and kraft lignin. These datasets are currently being used in a companion study to investigate common pathways for LBP metabolism in *R. palustris* (18). Using transcriptomics and proteomics data as input features and growth rates as the target, three machine learning models—Artificial Neural Networks (ANN), Random Forest (RF), and Support Vector Machines (SVM)—were trained, with the ANN model achieving the highest accuracy for both transcriptomic (94%) and proteomic (96%) datasets. Further analysis using permutation feature importance on the ANN models identified the top twenty genes and proteins influencing growth rates. By integrating the feature importance scores from both datasets, eight transport proteins were found to significantly impact *R. palustris* growth during LBP catabolism. Retraining the ANN model based on these eight transport proteins yielded prediction accuracies of 86% for proteomic data and 76% for transcriptomic data. Overall, this work demonstrates the utility of ANN models in identifying growth-associated genes and proteins that regulate the metabolic responses of *R. palustris* under aerobic and anaerobic conditions while catabolizing LBPs.

## Materials and Methods

### Growth experiments of *R. palustris* for different LBPs

The strain *Rhodopseudomonas palustris* BAA-98 CGA009 was sourced from the American Type Culture Collection (ATCC). All strains utilized in this study were preserved at −80°C. For storage, *R. palustris* strains were frozen in a 20% (v/v) glycerol solution, while *E. coli* strains were stored in 15% (v/v) glycerol. Upon recovery, *R. palustris* strains were cultured on solid 112 Van Niel’s medium, and E. coli on LB agar (Miller, AMRESCO), both supplemented with the required antibiotics (46).

For experimental setup, *R. palustris* seed cultures were first grown aerobically. Light-dependent biomass production (LBP) assays were conducted using 50 mL of Photosynthetic Medium (PM) in 250 mL Erlenmeyer flasks, enriched with 20 mM sodium acetate, 10 mM bicarbonate, and 15.2 mM ammonium sulfate. Aerobic seed cultures were diluted to an OD660 of 0.2 (1/10^th^ of their initial OD) and grown either in 50 mL PM in Erlenmeyer flasks or anaerobically in 13.5 mL PM within sealed 14 mL round-bottom tubes. The anaerobic cultures were exposed to light during growth. The light source used was an upper LED shelf light (SN-AG230-WIR-065) within an Algaetron incubator from Photon System Instruments (PSI). Both conditions were maintained at 30°C with shaking at 275 rpm. For all cultures, 10 mM bicarbonate, 15.2 mM ammonium sulfate, and either 1 mM or 10 mM LBP were added as needed. Each data point in the growth curve represents the average of three biological replicates.

### Transcriptomics data generation

Firstly, ribosomal RNA depletion was performed using NEBNext® rRNA Depletion Kit (Bacteria, New England Biolabs cat#E7860S). The rRNA-depleted RNA was purified by 2x RNAClean XP beads (Beckman Coulter) and eluted in 45ul of nuclease-free water. The purified RNA was then mixed with 4ul of NEBNext First Strand Synthesis Reaction Buffer and 1ul of random primers. The reaction was incubated at 94C for 12 mins for fragmentation. After incubation, the cultures were centrifuged to obtain cell pellets, which were then resuspended in RNAlater to prevent RNA degradation. Subsequently, to perform first strand cDNA synthesis, 8ul of NEBNext Strand Specificity Reagent and 2ul of NEBNext First Strand Synthesis Enzyme Mix was added to the reaction and the reaction was incubated at 25C 10 mins, 42C 30 mins, 70C 15 mins. Second strand reaction was then performed by adding 8ul of NEBNext Second Strand Synthesis Reaction Buffer with dUTP Mix (10X), 4ul of NEBNext Second Strand Synthesis Enzyme Mix, and 48ul of nuclease-free water. The reaction was incubated at 16C 1hr. The cDNA was purified by 1.8x SPRIselect Beads (Beckman Coulter) and eluted in 50ul of nuclease-free water. The reaction was incubated at 16C 1hr. The cDNA was purified by 1.8x SPRIselect Beads (Beckman Coulter) and eluted in 50ul of nuclease-free water. Subsequently, endprep reaction was performed by adding 7ul of NEBNext Ultra II End Prep Reaction Buffer and 3ul of NEBNext Ultra II End Prep Enzyme Mix into 50 ul purified cDNA. Endprep reaction was incubated at 20C 30 mins and 65C 20 mins. Adaptor ligation reaction was then performed by adding 1ul of NEBNext Ligation Enhancer, 30ul of NEBNext Ultra II Ligation Master Mix, and 2.5ul of NEBNext Adaptor, diluted to 0.5uM in Adaptor Dilution Buffer. The mix was incubated at 20C 15 mins. 3ul of USER Enzymer (New England Biolabs) was then added to the ligation product and the reaction was incubated at 37C 15 mins. The ligated product was purified by SPRIselect Beads (Beckman Coulter) and eluted in 15ul of nuclease-free water. PCR was carried out by adding 25ul of NEBNext Ultra II Q5 Master Mix, 5ul of i5 Primer, and 5ul of i7 Primer into 15ul of purified ligated product. PCR was performed at 98C 30 seconds, 15 cycles of 98C 10 seconds and 65C 75 seconds, and a final extension at 65C 5 mins. The final library was then purified by SPRIselect Beads and loaded into Illumina NovaSeq6000 paired end 150 bp mode for sequencing.

Data Quality Control After obtaining the raw data (fastq files), the quality of the original reads including sequencing error rate distribution and GC content distribution, is evaluated using FastQC. The original sequencing sequences contain low quality reads and adapter sequences. To ensure the quality of data analysis, raw reads must be filtered to get clean reads, and the subsequent analysis is based on clean reads. Data filtering mainly includes the removal of adapter sequences in the reads, the removal of reads with high proportion of N (N denotes the unascertained base information), and the removal of low-quality reads. This process is carried out using fastp.

To remove ribosomal RNA (rRNA) from bacterial transcriptomic data, we utilized the SortMeRNA tool, a specialized local sequence alignment program known for its efficiency in filtering rRNA from high-throughput sequencing reads. Installed via Conda for assured compatibility, SortMeRNA employs an approximate seed-based algorithm that enables swift and sensitive rRNA identification and separation from the transcriptomic data.

The application of SortMeRNA resulted in the segregation of rRNA reads from non-rRNA reads, providing us with a refined dataset focused on coding regions and other functional genes. This targeted removal of rRNA components is pivotal for subsequent transcriptome assembly and functional gene analysis, ensuring that our research delves into the biologically relevant aspects of the transcriptome unencumbered by the predominance of rRNA sequences. The reference genome index was created by the build-index function in HISAT2 software package with default options. Then the filtered clean reads are mapped to reference genome by HISAT2, the position and gene characteristics information were acquired. After the alignment, the generated SAM files were sorted to BAM files using samtools.

We used featureCounts software of subread package to quantify gene expression levels using mapped reads’ positional information on the gene. The genes number in different expression levels as well as the gene expression level of each single gene were analyzed. DESeq was used to analyze the DEG (differentially expressed genes) for samples with biological replicates and used edgeR for the samples without replicates. During the analysis, samples should be firstly grouped so that comparison between every two groups as a control-treatment pairwise can be done later. During the process, Fold Change ≥1.5 and FDR <0.05 are set as screening criteria. **<u>Proteomics data generation</u>**

The cell pellets were lysed in Pierce RIPA buffer (Thermo Fischer Scientific, Waltham, MA, USA) containing 5 mM DTT and 1x protease inhibitor (c*O*mplete EDTA-free protease inhibitor cocktail; Roche, Indianapolis, IN, USA) by shaking them at 95°C for 10 min on a thermomixer. The samples were then centrifuged at 16,000 x g for 15 min, and the supernatants were transferred to a new tube. The proteins were assayed using the CB-X protein assay (G-Biosciences, St Louis, MO, USA). 50 µg of reduced protein was alkylated with 20 mM iodoacetamide for 40 min and quenched with DTT. The proteins were then precipitated with acetone and the pellets washed 3 times with 70% ethanol. Proteins were resuspended in 50 µL of 50mM Tris/HCl, pH 8.0 containing 1 µg Lys-C and digested for 4 h, followed by further digestion with 1 µg trypsin overnight at 37°C. A quality control reference sample was prepared by mixing all the samples with the same 1:1 ratio to run between every 16 samples to check for instrument performance deviation. The sequence order of the samples was randomized using block randomization.

Each digest was run by nano liquid chromatography-tandem mass spectrometry (nanoLC-MS/MS) using an Ultimate 3000 RSLCnano system coupled to an Orbitrap Eclipse mass spectrometer (Thermo Fisher Scientific). Briefly, peptides were first trapped and washed on a trap column (Acclaim PepMap™ 100, 75µm x 2 cm, Thermo Fisher Scientific). Separation was then performed on a C18 nano column (Acquity UPLC® M-class, Peptide CSH™ 130A, 1.7µm 75µm x 250mm, Waters Corp, Milford, MA, USA) at 300 nL/min with a gradient from 5-22% over 75 min. The LC aqueous mobile phase was 0.1% (v/v) formic acid in water and the organic mobile phase was 0.1% (v/v) formic acid in 100% (v/v) acetonitrile. Mass spectra were acquired using the data-dependent mode with a mass range of m/z 375–1500, resolution 120,000, AGC (automatic gain control) target 4 x 10^5^, maximum injection time 50 ms for the MS1. Data-dependent MS2 spectra were acquired by HCD in the ion trap with a normalized collision energy (NCE) set at 30%, AGC target set to 5 x 10^4^ and a maximum injection time of 86 ms.

The identification and quantitation of the proteins were done using Proteome Discoverer (Version 2.4; Thermo Fisher Scientific) utilizing MASCOT search engine (Version 2.7.0; Matrix Science Ltd, London, UK). The search was performed against an in-house modified version of the cRAP database (theGPM.org/cRAP) and the *Rhodopseudomonas palustris* (version_20230110) database obtained from UniProt (ID: UP000001426_258594, www.uniprot.org), assuming the digestion enzyme trypsin and a maximum of 2 missed cleavages. Mascot was searched with a fragment ion mass tolerance of 0.06 Da and a parent ion tolerance of 15.0 ppm. Deamidated of asparagine and glutamine, oxidation of methionine, were specified in Mascot as variable modifications while carbamidomethylation of cysteine was fixed. Peptides were validated by Percolator with a 0.01 posterior error probability (PEP) threshold. The data were searched using a decoy database to set the false discovery rate (FDR) to 1% (high confidence). Only proteins identified with a minimum of 2 unique peptides and 5 peptide-spectrum matches (PSMs) were further analyzed for quantitative changes. The peptides were quantified using the precursor abundance based on intensity. The peak abundance was normalized using total peptide amount. The peptide group abundances are summed for each sample and the maximum sum for all files is determined. The normalization factor used is the factor of the sum of the sample and the maximum sum in all files. The normalized abundances were scaled for each protein so that the average abundance is 100.

### Different ML algorithms

Three distinct machine learning models—Artificial Neural Network (ANN), Random Forest (RF), and Support Vector Regression (SVR)—were trained to predict microbial growth rates using a high-dimensional proteomics dataset comprising 1854 features. The dataset was loaded into a pandas DataFrame, and feature standardization was performed using StandardScaler to ensure zero mean and unit variance, a critical preprocessing step when dealing with models sensitive to feature scaling. The ANN model was implemented as a deep Multi-Layer Perceptron (MLP) with 12 hidden layers, each containing 200 neurons, resulting in a highly parameterized model. The training process was carried out over 1000 iterations (epochs) using a random seed of 42 for reproducibility. To further interpret the model, permutation feature importance was computed over 30 iterations, identifying the top twenty proteins most predictive of the growth rate, revealing critical biological markers. The Random Forest (RF) regressor was implemented with 1000 estimators (decision trees), a model configuration designed to capture complex interactions between features while reducing variance through ensemble learning. The model’s inherent feature importance mechanism was used to assess the contribution of individual proteomic features, further aiding in the biological interpretation of the data. For the Support Vector Regression (SV) model, a radial basis function (RBF) kernel was selected to capture nonlinear relationships within the proteomic data. Hyperparameters were tuned with a regularization parameter (C) set to 100 and epsilon set to 0.01, optimizing the model’s balance between bias and variance.

### Statistical analysis for top genes and proteins

Statistical analyses on the nine statistically significantly abundant proteins (between aerobic and anaerobic conditions) to identify proteins with highest expression variability. For each protein of those nine proteins, the mean expression level and standard deviation were calculated across the fourteen conditions. To normalize the variability across proteins, the coefficient of variation (CV) was calculated as the ratio of the standard deviation to the mean expression level. The proteins were then ranked according to their CV values, with lower CV values indicating more consistent expression levels between aerobic and anaerobic conditions. Conversely, proteins with higher CV values exhibited greater variability between aerobic and anaerobic conditions.

### Simulation platform

ANN, RF, SV, and CV algorithms were implemented in Python using an Intel(R) Core(TM) i5-8250U CPU 1.60 GHz HP laptop with 8.00 GB of RAM and 64-bit operation with Windows 11 Home operating system.

## Supporting Information

Figure S1. Growth curve of *R. palustris* in different LBPs for aerobic and anaerobic conditions.

Figure S2. StringDB network of top twenty proteins.

Figure S3. Co-efficient of variation for top nine differentially abundant between aerobic and anaerobic conditions.

Table S1. Harvesting time for different conditions.

Table S2. Top twenty genes impacting the growth rates.

Table S3. Top twenty proteins impacting the growth rates.

Table S4. Combined list of top twenty proteins.

## Acknowledgements

R.S. gratefully acknowledges funding support from the National Science Foundation (NSF) CAREER grant (1943310).

## Authors Contributions

R.S. designed the study; oversaw the project and funding acquisition; N.B.C. performed data curation, formal analysis, validation, and visualization; N.B.C., M.K., and N.S. worked on methodology; N.B.C. wrote the original draft; all the authors reviewed and edited the draft.

## Data Availability

The data that support for the findings of this study can be found in the related cited articles and/or in the supplementary data. All the codes used to generate these results can be accessed in the GitHub repository (https://github.com/ssbio/r_palustris_LBP_ML).

