## Supplemental Figures for "Artificial Neural Network Reveals the Role of Transport Proteins in *Rhodopseudomonas palustris* CGA009 During Lignin Breakdown Product Catabolism"

Running Title: Artificial neural Network on *R. palustris* CGA009 "Omics" Data

### Supplementary Figures

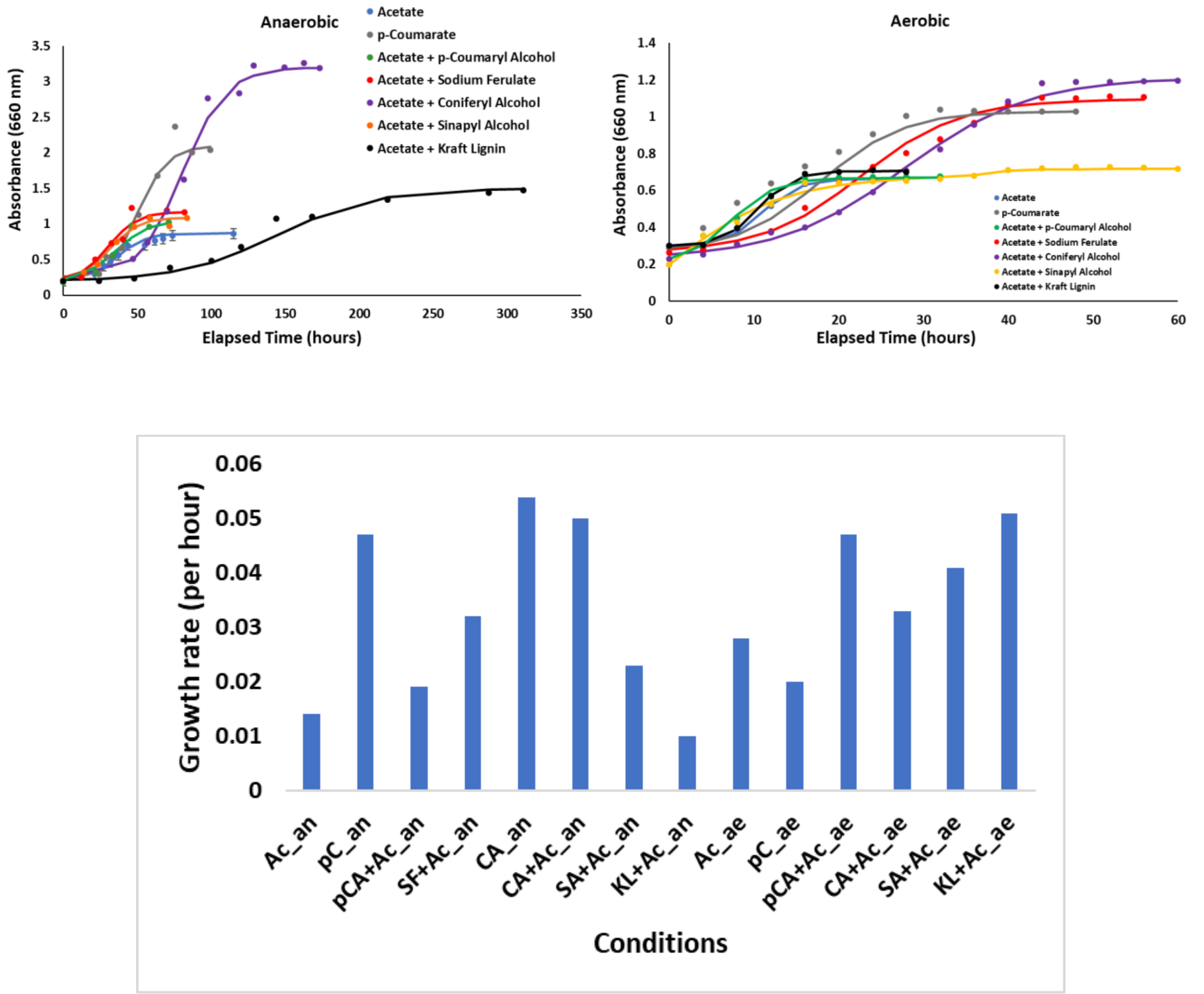


**Figure S1.** Growth curve of *R. palustris* in different LBPs for aerobic and anaerobic conditions. Here an indicates anaerobic and ae indicates aerobic. Further annotations for different substrates are Ac- acetate, pC- *p*-coumarate, pCA- *p*-coumaryl alcohol, SF- sodium ferulate, CA- coniferyl alcohol, SA- sinapyl alcohol, KL- kraft lignin.

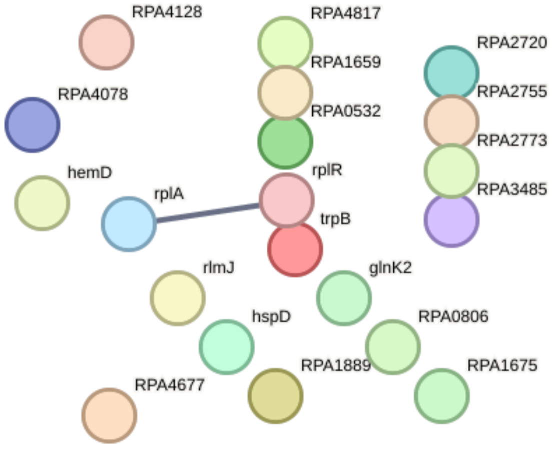


**Figure S2.** StringDB network of top twenty proteins.

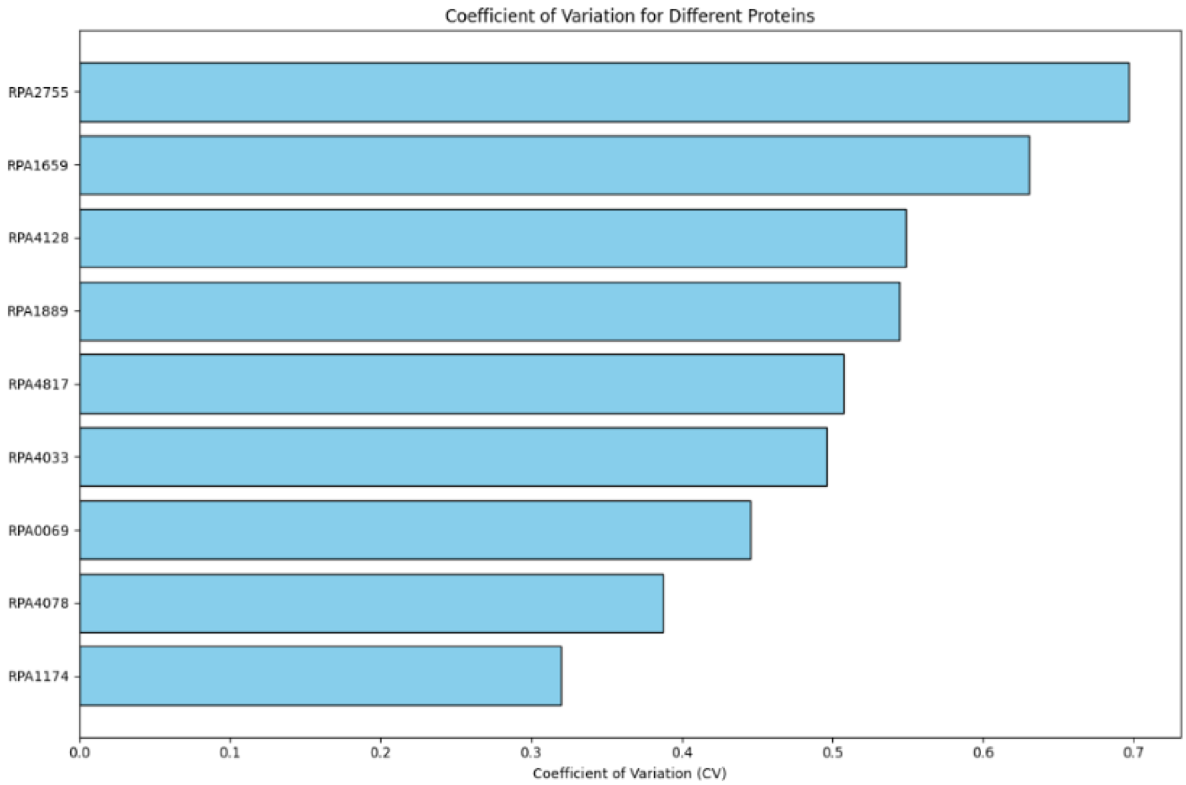


**Figure S3.** Co-efficient of variation for top nine differentially abundant between aerobic and anaerobic conditions.
